# Minimal RNA replicons for targeted gene silencing based on an asymptomatic viroid

**DOI:** 10.1101/2022.02.04.479109

**Authors:** Joan Marquez-Molins, María Urrutia-Perez, Andrea Gabriela Hernandez-Azurdia, Vicente Pallas, Gustavo Gomez

**Affiliations:** Institute for Integrative Systems Biology (I2SysBio), Consejo Superior de Investigaciones Científicas (CSIC) - Universitat de València (UV), Parc Científic, Cat. Agustín Escardino 9, 46980 Paterna, Spain; Instituto de Biología Molecular y Celular de Plantas (IBMCP), Consejo Superior de Investigaciones Científicas (CSIC) - Universitat Politècnica de València, CPI 8E, Av. de los Naranjos s/n, 46022 Valencia, Spain

**Keywords:** Viroid, gene silencing, RNA replicon, *Avsunviroidae*, eggplant

## Abstract

Gene silencing for functional studies in plants has been largely facilitated by manipulating viral genomes with inserts from host genes to trigger virus induced gene silencing (VIGS) against the corresponding mRNAs. However, viral genomes encode multiple proteins and disrupt plant homeostasis by interfering with endogenous cell mechanisms. To circumvent this issue, we have developed a silencing method based on the minimal autonomously-infectious nucleic acids currently known: viroids. In particular, *Eggplant latent viroid* (ELVd), an asymptomatic viroid, was manipulated with insertions between 21 to 42 nucleotides and our results show that larger insertions are tolerated but secondary structure is critical for their stability. Additionally, these ELVd constructs are able of local and systemic spread and can silence a target gene in eggplant. Inspired by the design of artificial microRNAs, we have developed a standardized procedure to generate stable insertions into the ELVd genome capable of silencing the desired target gene. Analogously to VIGS, we have termed our approach Viroid Induced Gene Silencing (VdIGS) and demonstrate that is a promising tool for dissecting gene functions in eggplant. Overall, this represents the use of minimal circular replicating RNAs able to spread systemically combined with the production of a tailored sRNA for targeted silencing.

## Introduction

Gene silencing for functional studies in plants has been largely facilitated by manipulating viral genomes with inserts from host genes to trigger virus induced gene silencing (VIGS) against the corresponding mRNAs (1, 2). This technology has been adapted for high-throughput functional genomics as it by-passes the labour extensive plant transformation (3). Therefore, transient silencing of endogenous plant genes, for basic research or biotechnological application, has been achieved using a wide variety of plant viruses (4). However, viral genomes encode multiple proteins and disrupt plant homeostasis by interfering with endogenous cell mechanisms (5).

To circumvent this issue, we have developed a silencing method based on the minimal autonomously-infectious nucleic acids currently known: viroids (6, 7). These RNA pathogens, naturally found infecting higher plants, are characterized by their extremely reduced genome size (246-401 nt) (8). Its mature form is a covalently closed, singlestranded RNA molecule with a highly compact structure which protects them against the RNA-silencing mediated degradation (9–11). According to the compartment in which replication takes place, viroids have been classified in two families: *Pospiviroidae* (nucleus) (12) and *Avsunviroidae* (chloroplast) (13). It has been described that viroids of both families trigger RNA silencing and thus viroid derived small RNAs (vd-sRNAs) are produced in infected plants (14–17), but it is unreported the use of any of these circular RNA replicons for specifically silencing a desired gene.

Although viroids can be pathogenic RNAs causing evident phenotypic alterations in host plants (18–20), completely asymptomatic viroid infections have also been reported, as is the case for eggplant latent viroid (ELVd) (21). This viroid was the latest member of the family *Avsunviroidae* formally characterized (22) and as the other members of this family replicates and accumulates in the chloroplast, although is also specifically trafficked to the nucleus (23). Interestingly, a mutational analysis of ELVd to inspect RNA processing led to the discovery that an eight-nucleotide insertion in the terminal loop of the right upper hairpin of ELVd was tolerated and partially maintained in the progeny, while all the other assayed mutations impede the infection of its natural host, eggplant (24).

Here, we have analyzed how larger insertions into this terminal loop affect ELVd viability and whether these insertions could cause the silencing of a target gene. Our results show that larger insertions are tolerated but secondary structure is critical for their stability into the ELVd genome. Additionally, these manipulated ELVd clones can efficiently target a desired gene in eggplant. Inspired by the design of artificial microRNAs (25, 26), we have developed a standardized cloning procedure into the ELVd genome for facilitating the silencing of a target gene. Analogously to VIGS, we have termed our approach Viroid Induced Gene Silencing (VdIGS) and demonstrate that is a promising tool for dissecting gene functions in eggplant.

## Material and methods

### Molecular cloning

The lethal gene *ccdB* flanked by *BsmBI* sites was inserted into the positions 245 and 246 of *Eggplant latent viroid* (ELVd) accesion AJ536613.1 and cloned into a pBluescript SK plasmid using high-fidelity PrimeSTAR HS DNA Polymerase (Takara, Kusatsu, Japan). This plasmid linearized with *BsmB*I was used to ligate self-hybridized oligonucleotides into the positions 245 and 246 of ELVd. The oligonucleotides were self-hybridized by denaturing 5 minutes at 95 °C and cooling down (0.05°C/sec) to 25°C and then ligated using T4 DNA ligase (Thermo Scientific™, Waltham, MA, USA). The resulting plasmids were used for generating the dimeric viroid sequences into a binary vector as it has been previously described (27). The oligonucleotides used to generate each insert are listed into the Supplementary table 1.

### RNA secondary structure predictions

ELVd (AJ536613.1) secondary structure of the plus polarity had been previously elucidated and the *in vivo* data obtained by high-throughput selective 2’-hydroxyl acylation analyzed by primer extension (hSHAPE) support the prediction of Mfold software (28). For our analysis, the minimal free energy was calculated considering that are circular molecules and a temperature of 25 °C degrees using Mfold (29). RNA secondary structures files were generated in Vienna format and were displayed using Forna (30).

### Viroid inoculation

Cotyledons of *Solanum melongena* cv Black beauty were inoculated by agro-infiltration with a culture of *Agrobacterium tumefaciens* strain C58 harbouring the pMD201t2 binary vector with the correspondent viroid-dimer. The overnight grown bacterial culture was diluted in infiltration buffer (MES 0.1 M, MgCl_2_ 1 M) up to an optical density at 600 nm of 1 and injected on the abaxial side of one cotyledon using a needle-less syringe. Plants were kept in a photoperiod of 16 hours under visible light and 25 °C (light)/22 °C (darkness). Local tissue was collected from six agro-infiltrated cotyledons to constitute a sample, while for systemic the upper leaf from at least three bio-replicates was taken.

### RNA extraction and northern/dot blot

Total RNA was extracted from systemic leaves using TRIzol reagent (Invitrogen, Carlsbad, CA, USA). For northern blot analysis 3 μg of total RNA per sample were mixed with solid urea and then loaded into a PAGE 5% urea 8 M and TBE 89 mM gel. RNA electrophoresis was performed at 200 V for 1 h and then RNA was transferred to a nylon membrane using MiniProtean 3 system (BioRad, USA). Transfer conditions were 100 V for 1 h at 4 °C in TBE buffer 1 ×. Nucleic acids transferred to the membrane were covalently fixed by using ultraviolet light (700 × 100 J/cm^2^). Dot-blot samples were directly applied into the membrane. Hybridization and chemiluminescent detection were performed as previously described (31).

### Small RNA purification and stem-loop RT

Low-molecular weight RNA (< 200 nt) were isolated from total RNA using REALTOTAL microRNA Kit RBMER14 (Durviz, Spain) according to the manufacturer’s instructions. Stem-loop-specific reverse transcription for small RNA detection was performed as previously described (32).

### RT-qPCR

First-strand cDNA was synthesized by reverse transcription using RevertAid cDNA Synthesis Kit (Thermo Scientific, USA). qRT-PCR assays were performed using PyroTaq EvaGreen mix Plus (ROX) (CulteK Molecular Bioline, Spain) according to the manufacturer’s instructions. All analyses were performed in triplicate on an ABI 7500 Fast-Real Time qPCR instrument (Applied Biosystems, USA) as previously described (33). The efficiency of PCR amplification was derived from a standard curve generated by four 10-fold serial dilution points of cDNA obtained from a mix of all the samples. The sequence of all primers is in Supplemental Table 1. Relative RNA expression was determined by using the comparative ΔΔCT method (34) and normalized to the geometric mean of cyclophilin (SMEL_001g116150.1.01) and F-Box (SMEL_002g166030.1.01), as references for mRNA whereas the small nucleolar U6 RNA (SMEL3Ch05) was used for normalizing sRNA-quantitation. The statistical significance of the observed differences was evaluated by a paired t-Test.

## Results

### Secondary structure is critical for the stability of insertions into the viroid genome

In order to test whether larger insertions, with the potential to generate small RNAs specific to a target gene, could be relatively stable into the ELVd genome (Figure 1A), several constructs were obtained. Firstly, 21-, 27-, 32- and 42-nucleotide insertions into the positions 245 and 246 of the wild type ELVd genome accession AJ536613.1 (Figure 1A, indicated in orange) were obtained (Figure 1B). These constructs were assayed and their expression in the infiltrated tissue was confirmed at 2 two days post inoculation (dpi) (Figure 1C). Northern blot analysis (Figure 1D) revealed that the ELVd-chimeras with the smaller insertions (21- and 27-nucleotides) failed to form the circular mature forms associated to viroid replication in local tissues, indicating that those secondary structures made impossible the transcription and/or circularization of the chimeric viroid. Conversely, circular mature forms carrying the 32- and 42-insertions were detected in local (2 dpi) and also systemic tissue (6 dpi) (Figure 1D, upper and lower panels, respectively). It is worth noting that the terminal loop of the right upper hairpin is more enlarged in the 21- and 27-nt insertions than in the 32- and 42-nt insertions which form a hairpin with an internal small loop. To validate the hypothesis than an enlarged loop compromises the viability of the chimeric ELVd, three new 42-nt insertions that enlarged the loop in one to three nucleotides were assayed (Supplementary Figure S1). The results show that mature circular forms were detected at 6 dpi for the constructs with one and two extra unpaired nucleotides, but not for the construct with three additional unpaired nucleotides in the loop.

**Figure 1.**
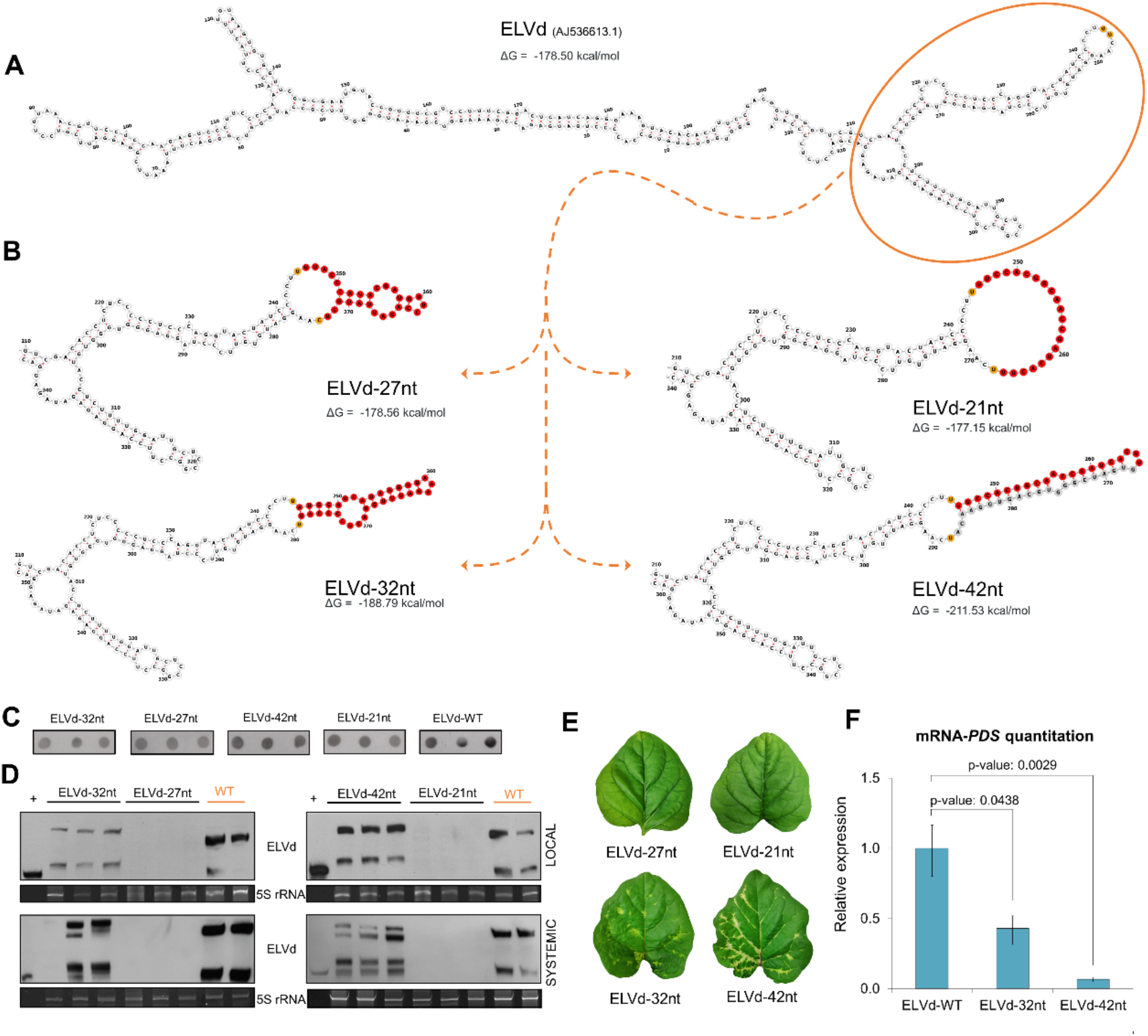
Stability of insertions in ELVd genome and induced silencing of the phytoene desaturase (PDS). (**A**) Secondary structure predictions with mfold at 25 °C displayed with forna of ELVd accession AJ536613.1 and its minimal free energy. (**B**) Secondary structure predictions with mfold at 25 °C displayed with forna of the insertions (in red) into the ELVd genome and the minimal free energies of the complete structures. In the case of the 42-nucleotide insertion, the nucleotides designed to target the PDS are depicted in red, while the complementary nucleotides necessary to form a hairpin-like structure are in grey (**C**) Dot blot hybridization of RNA from in infiltrated tissue at two days post inoculation (dpi). (**D**) Northern blot of RNA from local (6 dpi) and systemic (25 dpi) tissue. (**E**) Phenotype of the upper leaf at 25 dpi. (**F**) RT-qPCR relative accumulation of the mRNA of PDS for ELVd-WT, ELVd-32nt and ELVd-42nt. The statistical significance was estimated by paired T-tests and the obtained p-values are shown. Error bars represent the standard error values.

### Silencing of phytoene desaturase mediated by chimeric ELVd

These results indicate than ELVd-chimeras with insertions representing more than 10% of the genome (333nt) can replicate and move to distal parts of the plant as long as the secondary structure of the viroid is not compromised. The objective of these insertions was to target the phytoene desaturase (*PDS*) gene, a recognized indicator gene for VIGS, which encodes an enzyme that catalyzes an important step in the carotenoid biosynthesis pathway, resulting in a characteristic photo-bleaching when the gene is silenced. The hairpin of the 32-nt insertion is present in the *PDS* mRNA, while the 42-nt insertion is formed by the unstable 21-nt (Figure 1B upright), which were designed to target the *PDS*, and 21 additional nucleotides to form a hairpin. Visual inspection of systemic leaves at 4 weeks post inoculation (wpi) showed the characteristic whitening attributed to *PDS* silencing (Figure 1E), which was more intense in the ones infected with the ELVd carrying the 42-nt insertion. This was confirmed by RT-qPCR quantitation of the *PDS* mRNA (Figure 1F).

Although the 32-nt natural PDS hairpin was able to induce the silencing of the corresponding mRNA, we envisioned that it was more appealing to propose a strategy for Viroid Induced Gene Silencing (VdIGS) independent of the secondary structure of the target mRNA, which was additionally supported by the higher silencing observed (Figure 1F).

Consequently, for further characterization of the *PDS* silencing, we used the construct with the 42-nt insertion in which half of the insertion is designed for targeting the *PDS* and the other half for structural compensation (from now on termed as ELVd-PDS). PDS-silencing phenotype was observed since 3 wpi and at 25 dpi all plants inoculated with ELVd-PDS displayed whitening in systemic leaves, in contrast to the asymptomatic mock plants infected with the wild-type (WT) (Figure 2A). Northern blot analysis at four time points (23, 25, 27 and 29 dpi) detected the ELVd-PDS chimera (375 nt) but also the reversion to WT (333 nt) (Figure 2B). The maintenance of the 42-nt insertion was further confirmed by sequencing (Supplementary Figure S2). ELVd-PDS was the majoritarian form at 23 and 25 dpi and it decreased in favour to the WT version over time. To analyse up to what extent the chimeric ELVd persisted in the infected plants, despite its obvious lower fitness in comparison with the WT, we sampled 10 plants at two months post inoculation (Supplementary Figure S3). As expected, ELVd-WT was the predominant form, but the chimeric ELVd-PDS was also detected in most plants but in a much lower concentration (Supplementary Figure S3A). In fact, whitening in leaves was still observed in 9/10 plants (Supplementary Figure S3B). Overall, these results demonstrate that the chimeric ELVd-PDS is relatively stable in infected plants and if present produces the whitening associated to *PDS*-silencing.

**Figure 2.**
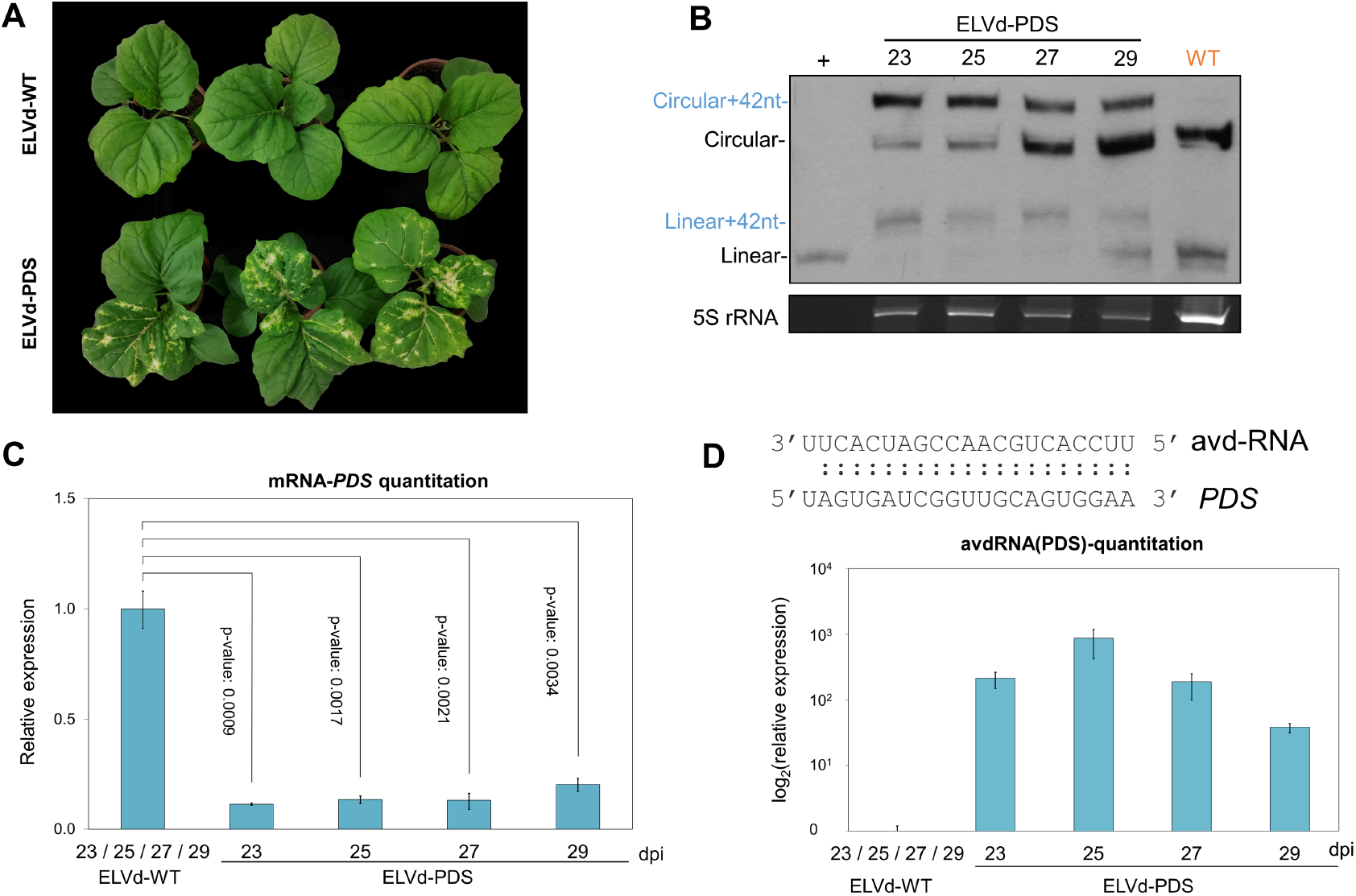
Silencing of the phytoene desaturase (PDS) induced by an artificial viroid derived sRNA. (**A**) Phenotype of plants infected with ELVd-WT (upper) and ELVd-PDS (lower) at 25 days post inoculation (dpi). (**B**) Northern blot of RNA from plants infected with ELVd-PDS at 23- 25- 27- and 29- dpi and an RNA mix of plants infected with ELVd-WT at these same timepoints. (**D**) RT-qPCR relative accumulation of the PDS mRNA for ELVd-WT and ELVd-PDS at 23- 25- 27- and 29- dpi. The statistical significance was estimated by paired T-tests and the obtained p-values are shown. (**D**) Quantitation of the artificial viroid-derived small RNA (avd-RNA) designed to silence the PDS mRNA in the infected plants by stem-loop RT-qPCR. The sequence of the avd-RNA and the target region in the mRNA of PDS are indicated above. Error bars represent the standard error values.

In order to efficiently target the PDS, the construct ELVd-PDS contains a sRNA of 21-nt designed in an analogous fashion as artificial microRNAs to cleave the *PDS* mRNA (see material and methods for details) and thus we denominated it artificial viroid-derived small RNA (avd-sRNA). To validate that this avd-sRNA designed to target the PDS was indeed being produced, stem-loop RT-qPCR was used to quantify its accumulation in the four timepoints mentioned above (Figure 2C). As expected, this avd-sRNA was not detected in mock samples. The levels of avd-sRNA peaked at 25 dpi and subsequently decreased, which can be explained according to the drop in the band intensity of the chimeric ELVd-PDS in the next timepoints (Figure 2B). However, *PDS* silencing quantified by RT-qPCR showed similar levels of decrease at 23, 25 and 27 dpi while at 29 dpi the *PDS* gene was slightly less silenced (Figure 2D). This shows that in this time-range the target gene is efficiently silenced.

### Amplification-free cloning of fragments into ELVd

We have designed a workflow for Viroid Induced Gene Silencing (VdIGS) in which a 21-nucleotide sequence (designed to silence a desired target gene) is structurally compensated by other 21-nt to form a hairpin (Figure 2). In this way, any 21-nucleotide insertion for targeting a certain gene, can be structurally compensated. The design of the 21-nucleotide sequence specific for silencing the target gene is based on that of artificial microRNAs (amiRNAs) (25, 26). Specifically, the software P-SAMS is used to design a 21-nt amiRNA and the star sequence (amiRNA*) that provides P-SAMS is used to form the hairpin (35). For cloning into the ELVd genome the resultant 42-nt (amiRNA-amiRNA*), self-hybridizing DNA oligonucleotides are designed with CCTT and TTGA overhangs as represented in Figure 2A. Therefore, the hybridized oligonucleotides can be directly ligated in the correct orientation into a plasmid harbouring the ELVd genome cDNA, which is further dimerized into a binary vector to obtain the infectious clone vector as previously described (27, 36).

### Assessment of the applicability of viroid induced gene silencing (VdIGS)

To evaluate whether, irrespectively of sequence, the aforementioned strategy can produce stable chimeric-ELVd that induce silencing, we obtained two constructs targeting another indicator gene, the magnesium-chelatase subunit *ChlI*. These constructs were designed as described in Figure 3A, and the insertions folded forming the expected hairpins (Supplementary Figure S4). The stability of the constructs was assessed in a northern blot at 25 dpi (Figure 3B). In both constructs the characteristic yellowing produced by the *ChlI* silencing was observed (Figure 3C). Additionally, this was corroborated by a significant decrease in the expression of this gene measured by RT-qPCR (Figure 3D). Therefore, these results prove that VdIGS is an effective technology for silencing a candidate gene.

**Figure 3.**
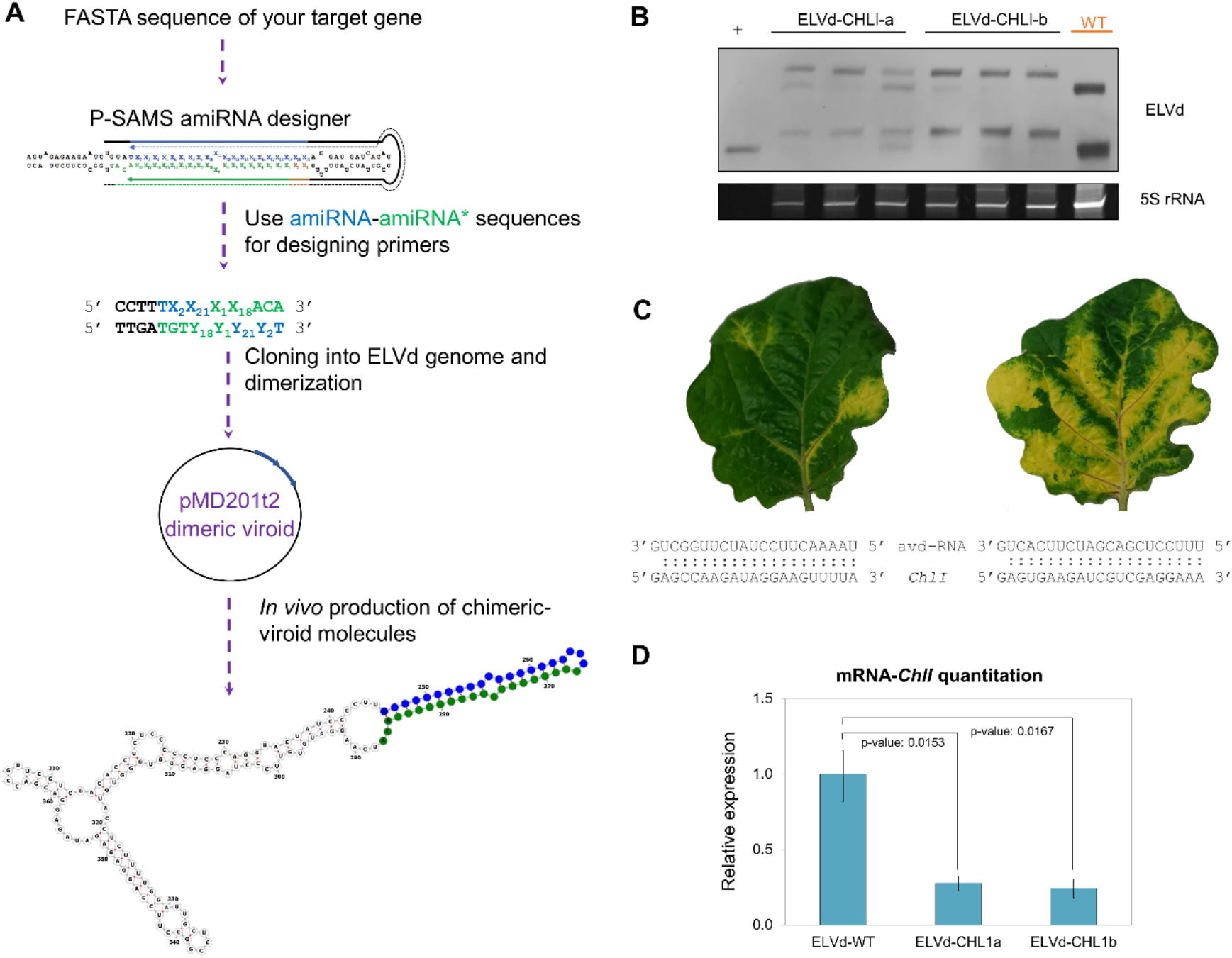
Viroid induced gene silencing (VdIGS)-Workflow for generating VdIGS constructs and assessment of the applicability. (**A**)To design a construct for targeting a desired gene, its sequence is introduced in the P-SAMS website for artificial microRNA (amiRNA) design. A pair consisting of an amiRNA (blue) and its star sequence (amiRNA*, in green) should be chosen. Subsequently, oligonucleotides with these sequences plus four nucleotides of the viroid (in black) are designed as represented. These oligonucleotides are self-hybridized and directly cloned into a plasmid harbouring the ELVd genome, in which the *ccdB* gene is excised by BsmbI digestion. Once obtained this construct, a dimer is generated into the binary vector pMD201t2 as previously reported (27). This plasmid is transformed into agrobacterium and the VdIGS RNAs are generated *in vivo* by transient expression. (**B**) Northern blot of RNA from plants infected with the constructs ELVd-CHLI-a and ELVd-CHLI-b at 25 days post-inoculation. (**C**) Phenotype of the upper leaf at 25 dpi of plants inoculated with ELVd-CHLI-a (left) and ELVd-CHLI-b (right). The sequence of the avd-RNA and the target in the mRNA of the magnesium-chelatase subunit ChlI are indicated below. (**D**) RT-qPCR relative accumulation of the mRNA of ChlI. The statistical significance was estimated by paired T-tests and the obtained p-values are shown. Error bars represent the standard error values.

## Discussion

In this work we have used insertions into an asymptomatic viroid to study the stability and the potential use of constructs harbouring these insertions to induce gene silencing. Interestingly, we found that the secondary structure was more crucial than the insert size for permitting viroid replication and/ or circularization. Moreover, our results show that artificial viroid derived sRNAs (avd-sRNAs) can efficiently target a desired gene in eggplant. This technique combines the target specificity of 21 nt small RNAs and the systemic movement of a viroid, being a novel tool for plant biotechnology.

Thus, VdIGS can be used for studying gene-function using an asymptomatic RNA replicon in eggplant, which is among the top five most important vegetables in the Mediterranean and Asia (37) and its genome has been recently re-sequenced (38–40) evidencing that it is a well-studied crop with on-going attention. Another advantage is the cloning procedure which does not requires the amplification of the target gene as in VIGS (typical insert size is 300 nt) because oligonucleotides (46-nt in length) are self-hybridized and cloned into the ELVd genome.

Additionally, our results might have implications for understanding viroid biology and evolution. We show that an enlarged loop abolished the accumulation of mature forms and only hairpin structures were stable. That observation explains why inserts with roughly the same size (40 nt) but with 16 nt folding into a loop failed to infect eggplant (41). Moreover, chimeric ELVd reverted to the wild-type over time, but no intermediate forms were detected, reinforcing the notion that the maintenance of a compact structure is essential for this viroid, and partial deletions of the artificial hairpins are not tolerated. The incorporation of novel sequences into the viroid genome is thus constrained by important structural limitations, and the insertions must not impede the transcription, circularization, and movement of these RNA replicons. Nonetheless, recombination is thought to play an important role in the evolution of viroids (42–44), and in fact, has been proposed as the only possible route to evolutionary innovation in these circular RNAs (11). The origin of viroids itself is a matter of considerable debate between the hypothesis of being ancient remnants of an RNA world (45, 46) or more recently escaped from a cellular context (47, 48). In any case, primordial RNA self-replicons should have some viroid-like features, and therefore understanding this intriguing RNAs may help learning about the origin of life (49). On the other hand, our results together with the association of peach calico to vd-sRNAs of *Peach latent mosaic viroid* (50–52) unarguably show that the small RNAs produced from chloroplast-replicating viroids can be functional. However, the biogenesis and DICER-loading of such vd-sRNAs is still unclear due to the absence of silencing machinery in this organelle and lack of known mechanisms of RNA transport. Further research will be required to clarify this issue.

In conclusion, here we have presented the use of minimal circular replicating RNAs able to systemic spread combined with the production of a tailored sRNA for targeted silencing. This is a conceptual novelty that further expands the biotechnological applications of viroids in general, and ELVd in particular (41, 53).

## ACKNOWLEDGMENTS

This work was supported by the Spanish Ministry of Economy and Competitiveness (co-supported by FEDER) Grants PID2019-104126RB-I00 (GG) and PID2020-115571RB-I00 (VP). J.M.M. was recipient of a pre-doctoral contract ACIF-2017-114 from the Generalitat Valenciana. The funders had no role in the experiment design, data analysis, decision to publish, or preparation of the manuscript.

